# Higher-order discrimination learning by honey bees in a virtual environment

**DOI:** 10.1101/403956

**Authors:** Alexis Buatois, Lou Laroche, Aurore Avarguès-Weber, Martin Giurfa

**Affiliations:** Research Centre on Animal Cognition, Center for Integrative Biology, CNRS, University of Toulouse, 118 route de Narbonne, F-31062 Toulouse cedex 09, France

**Author notes:** First authorship shared (Alexis Buatois, Lou Laroche). Senior authorship shared (Aurore Avarguès-Weber, Martin Giurfa). **Corresponding authors:** Drs. Martin Giurfa & Aurore Avarguès-Weber.

**Keywords:** Learning, Cognition, Non-elemental learning, Negative Patterning, Vision, Virtual Reality, Honey Bees

## Abstract

Non-elemental learning constitutes a cognitive challenge because, contrary to elemental learning forms, it does not rely on simple associations, as events to be learned are usually ambiguous in terms of reinforcement outcome. Negative patterning constitutes a paradigmatic case of non-elemental learning, as subjects have to learn that single elements A and B are reinforced while their conjunctive representation AB is not reinforced (A+, B+ vs. AB-). Solving this problem requires treating the compound AB as being different from the linear sum of its components in order to overcome stimulus ambiguity (A+/A- and B+/B-). The honey bee is the only insect capable of mastering negative patterning as shown by numerous studies restricted mainly to the olfactory domain. Here we studied the capacity of bees to solve a negative patterning discrimination in the visual domain and used to this end a virtual reality (VR) environment in which a tethered bee walking stationary on a treadmill faces visual stimuli projected on a semicircular screen. Stimuli are updated by the bee’s movements, thus creating an immersive environment. Bees were trained to discriminate single-colored gratings rewarded with sucrose solution (blue, green; A+, B+) from a non-rewarded composite grating (blue-green, AB-). Bees learned this discrimination in the VR environment and inhibited to this end linear processing of the composite grating, which otherwise is treated as the sum of its components. Our results show for the first time mastering of a non-linear visual discrimination in a VR environment by honey bees, thus highlighting the value of VR for the study of cognition in insects.

## Introduction

Learning is a fundamental capacity for individual survival as it renders a complex environment predictable. Learning strategies vary depending on the complexity of the problem to be solved. On the one hand, elemental forms of learning rely on acquiring simple associative links between events in animal’s world. A typical example is Pavlovian conditioning^1^, in which subjects learn a simple associative link between a neutral stimulus (conditioned stimulus, CS) and a biologically relevant stimulus (unconditioned stimulus, US). On the other hand, non-elemental learning does not rely on such simple associative links, as events to be learned are ambiguous because they are as often reinforced as non-reinforced^2-5^. A good example of this situation is the so-called negative-patterning discrimination^6-8^ in which subjects have to discriminate a non-reinforced conjunction of two elements A and B from its reinforced elements (i.e. AB– vs. A+ and B+)^9-10^. The ambiguity of the task resides in the fact that each element (A and B) is as often reinforced (when presented alone) as non-reinforced (when presented as a compound). To solve the problem, it is necessary to abolish spontaneous linear summation predicting that AB is twice as reinforced as the reinforced A and B^9^.

In mammals, elemental and non-elemental learning forms are mediated by different brain structures, thereby supporting the notion that they represent different levels of complexity. Specifically, the hippocampus is dispensable for learning elemental associations^3, 5^ but is required for certain forms of non-elemental learning involving conjunctive representations such as negative patterning, spatial learning or contextual fear conditioning^3, 5, 7, 11-18^. Yet, solving non-elemental discriminations is not a prerogative of vertebrates. Several forms of non-elemental learning have been shown in the honey bee, an insect with impressive learning capabilities^2, 19-20^. Negative-patterning solving has been shown in harnessed bees using the olfactory conditioning of the proboscis extension response (PER) ^21-23^. In this Pavlovian protocol, harnessed bees learn to associate an odorant as CS with a drop of sucrose solution as US; after successful learning they exhibit PER to the odorant predicting the food. In the negative patterning variant of this protocol, bees learn simultaneously to respond to odorants A and B (A+, B+) and to inhibit their response to the conjunction of both odorants (AB-)^24-29^. Similarly, to the vertebrate case, specific circuits of a higher-order brain structure, the mushroom bodies, are required to solve this task: blocking synaptic transmission at the level of these circuits suppresses the capacity to solve negative patterning but leaves intact the capacity to solve linear discriminations^29^.

Most of the higher-order learning phenomena found in bees have been shown in the context of training free-flying bees to collect sucrose solution associated with visual stimuli in mazes and other setups^20, 30-31^. Yet, non-linear visual discriminations have been scarcely studied in this context. In a single study, free-flying bees were trained to discriminate a yellow and a violet checkerboard, both rewarded, from a non-rewarded violet and yellow checkerboard, thus reproducing the logic of a negative patterning problem^32^. Although bees succeeded in this task, whether in other conditions they treated spontaneously the dual checkerboard as the simple sum of its components was not determined. Furthermore, the use of free-flying bees precluded a full control of the animal behavior (such as in PER experiments) and the coupling with invasive methods aimed at unravelling the mechanisms of this performance.

The use of virtual reality (VR) offers new possibilities to fill both voids. Recently, we established a VR environment in which bees walking stationary on a treadmill learn simple visual discriminations (differential conditioning: A+ vs. B-) of visual targets projected on a semicircular screen placed in front of them^33-34^. Experiments are performed under closed-loop conditions^34^ so that the images perceived by the bee are constantly updated by its movements, thus creating a sensation of immersion within this virtual environment. Although VR allows learning simple associations^33-35^, it might affect negatively the solving of non-linear problems because of the constraints it imposes on active vision^34^.

Here we studied the capacity of tethered bees to solve a negative-patterning discrimination in a VR context. We first determined that bees treat spontaneously a reinforced visual compound as the linear sum of its components. We then showed that when trained to do so, bees learn to inhibit this lineal processing to solve negative patterning under VR conditions. Our results show for the first time that bees master a non-linear visual discrimination in a VR environment, thus highlighting the value of VR for the study of the mechanisms underlying this performance.

## Materials

### Animal preparation

Honey bee foragers (*Apis mellifera*) were caught upon landing on a gravity feeder and before they started collecting sucrose. They were anesthetized on ice for 3 min in the laboratory. The wings were then cut and the thorax shaved to attach a tether on the thorax using UV cured dentine^34^. Once attached, bees were fed with 4μl of 0.9M sucrose solution and placed on a miniature treadmill during 3h to familiarize them with the tethering and the treadmill and to increase feeding motivation. Bees were then placed on the treadmill associated with the VR setup 1 min before the start of the training procedure. Details on the VR setup (Fig.1a) are available in^34^ and in the electronic supplementary section.

**Figure 1.**
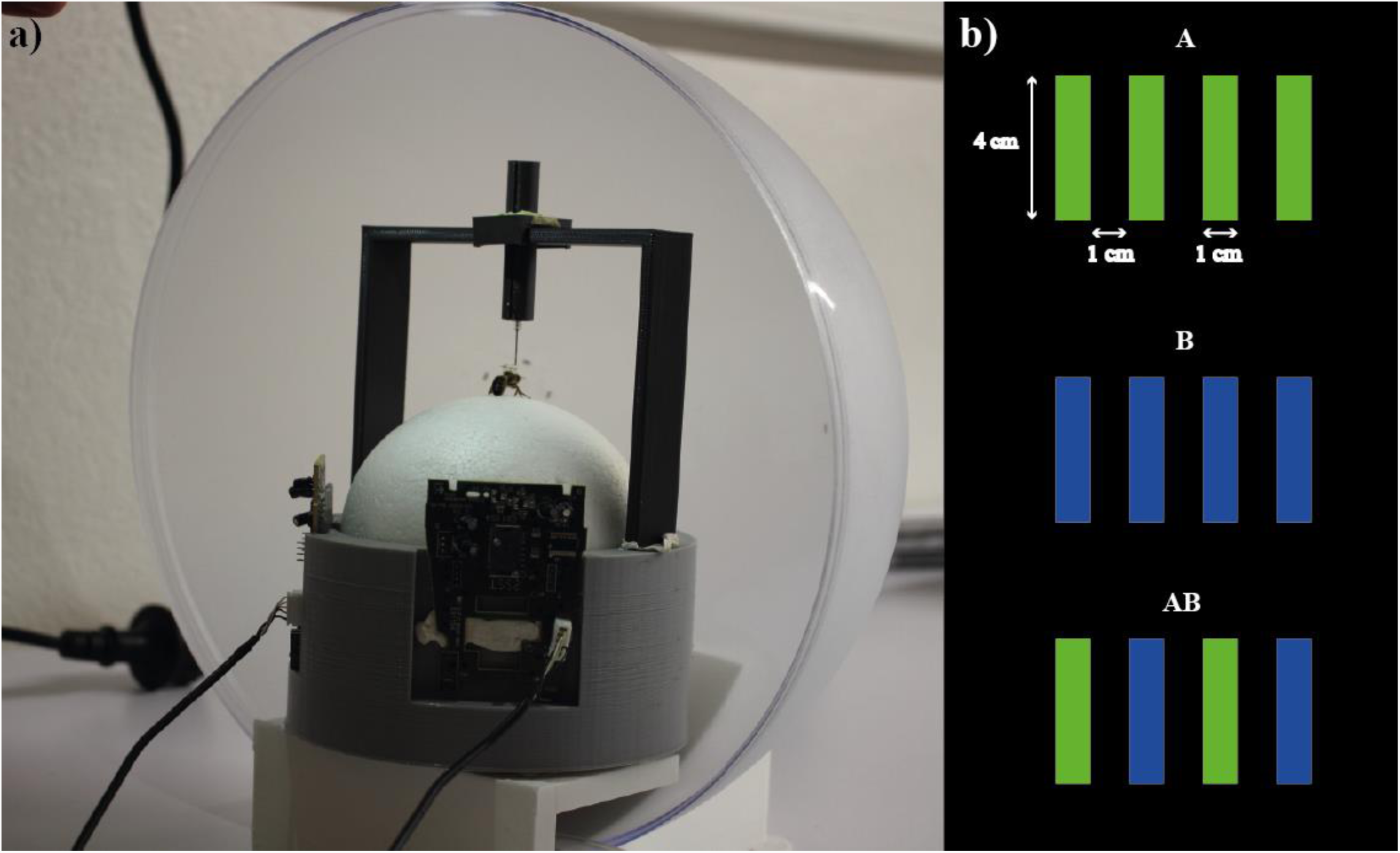
Virtual reality setup and visual stimuli. **(a)** Global view of the virtual reality system. **(b)** Conditioned stimuli: blue grating (A), green grating (B) and composite blue-green grating (AB).

### Visual stimuli

Visual stimuli (Fig.1b, Fig. S1, Table S1) were a green grating “A” (RGB: 0, 100, 0; irradiance = 24 370 μW.cm2; dominant wavelength = 530nm), a blue grating “B” (RGB: 0, 0, 255; irradiance = 161 000 μW.cm2; dominant wavelength = 450 nm), and a composite grating “AB” made of the two previous gratings (blue/green grating, irradiance = 116 347 μW.cm2). The irradiance of green bars was lowered with respect to that of the blue bars to reduce spontaneous attraction of naïve bees^34, 36^. Gratings were composed of four stripes, each measuring 1 cm width and 4 cm height. As the bee was placed at 10 cm from the displaying screen, each bar subtended a visual angle of 5.7° in the horizontal plane.

### Experiment 1: do bees treat a visual compound as the sum of its components?

Bees were trained to associate gratings A, B or AB (an independent group of bees for each training, n = 20 in each case; Table 1) with 1 M sucrose solution following an absolute conditioning regime (a single stimulus rewarded during training; see Fig. 2, left).

**Table 1:** Groups trained under an elemental-conditioning regime. Each group was trained with a different grating paired with sucrose solution. Following ten conditioning trials, bees were tested in the absence of reward in two dual choice situations opposing the grating previously trained and the two alternative gratings.

| Training Grating (CS+) | Test |
| --- | --- |
| Green+<br>(A) | Green vs. Blue; Green vs. Green-Blue<br>(A vs. B; A vs. AB) |
| Blue+<br>(B) | Blue vs. Green; Blue vs. Green-Blue<br>(B vs. A; B vs. AB) |
| Green-Blue+<br>(AB) | Green-Blue vs. Green, Green-Blue vs. Blue<br>(AB vs. A; AB vs. B) |

**Figure 2.**
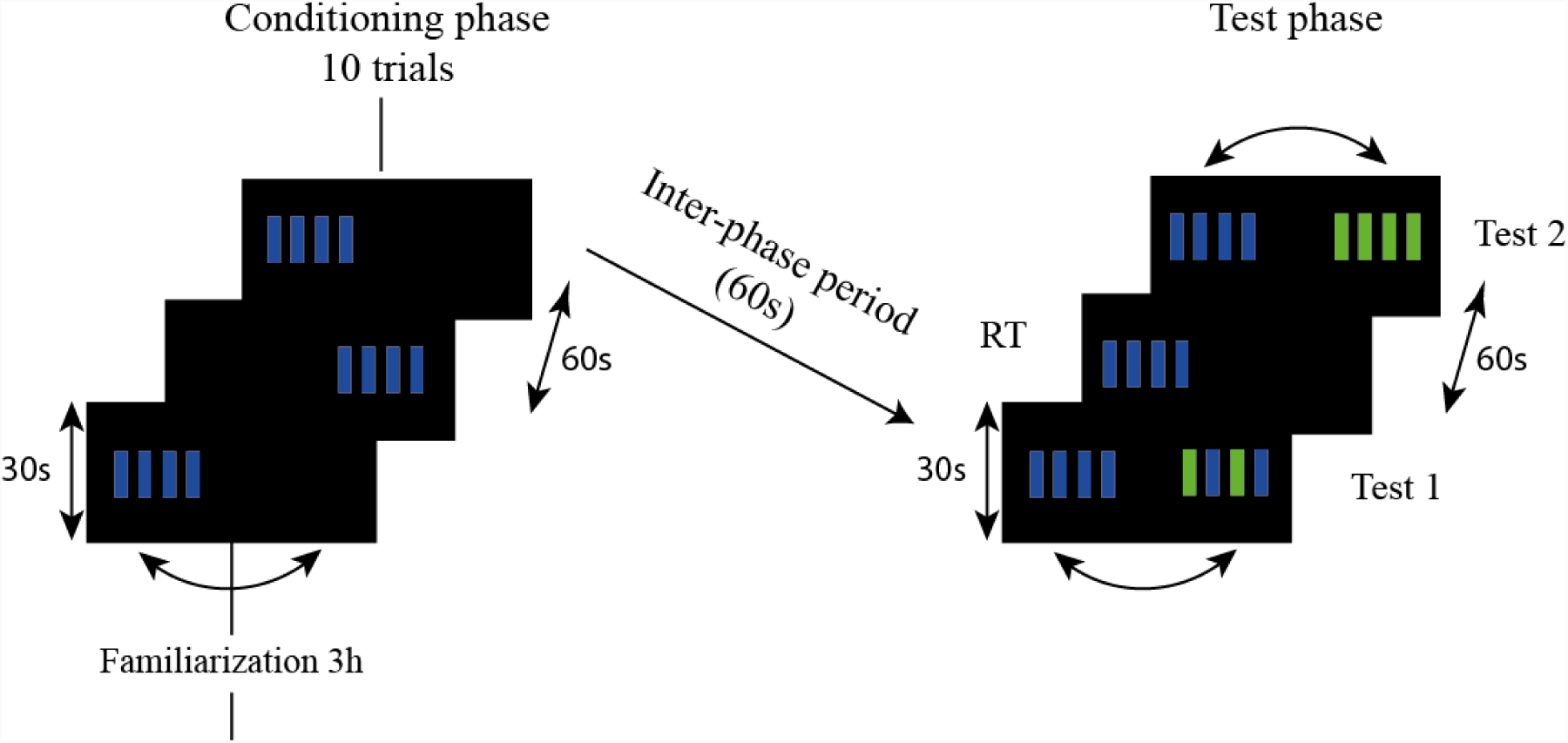
Experimental schedule of Experiment 1, illustrated in the case of blue-grating conditioning (B+). Bees started with a familiarization phase with the setup in the absence of visual stimulation, which lasted 3 hours. They were afterwards conditioned during 10 trials to associate the blue grating (B+) with a sucrose solution. The side of stimulus presentation was pseudo-randomized between trials. The grating was blocked during 8 s and rewarded with sucrose solution when bees aligned it with their body axis (i.e. when they centered it on the screen). Trials lasted a maximum of 30 seconds. They were ended after reward delivery. The intertrial interval was 60 s. After the end of conditioning, the bees were subjected to non-reinforced tests in which the conditioned stimulus (CS) was opposed to new stimuli (NS). In this example, the blue grating B was the CS and was therefore opposed to AB during the first test and to A during the second test. Test sequence was randomized from bee to bee. A refreshment trial was interspersed between tests to avoid extinction. The same schedule was followed in the case of green grating (A+) or composite green-blue grating (AB+) conditioning.

The conditioning phase lasted 10 trials (Fig. 2, left). At the beginning of each trial, the conditioned stimulus (CS) was presented to the right or left of the bee, following a pseudo randomized sequence (L, R, R, L, R, L, L, R, L, R; ± 50° from its body axis). When the bee aligned the CS with its body axis (0°), the stimulus remained in this position during 8 s and sucrose solution was first delivered to the antennae and then to the proboscis during 5 s by means of a toothpick. A trial lasted a maximum of 30 s and reward delivery set the end of the trial, even if the 30 s were not elapsed. A black background appeared then during 60 s before the start of a new trial. If the bee did not center the stimulus, the trial was ended after 30 s. No reinforcement was delivered in this case. This situation was infrequent due to phototaxis^33^. In average, a trial lasted 20 ± 3 seconds (mean ± S.E.)

One minute after the end of training, bees were tested in the absence of reward in two dual-choice situations opposing the CS and the two alternative stimuli (Table 1; Fig. 2, right). Each test lasted 30 s. At the beginning of each test, stimuli were placed randomly either to the right or the left of the bee (± 50° from its body axis). Centering the stimuli during these dual-choice situations did not result in blocking stimulus position in front of the bee. Thus, bees were able to switch from one stimulus to the other during the tests. Test sequence was randomized between bees. A refreshment trial lasting a maximum of 30 s was interspersed between the two tests. In this case, the CS was again rewarded to avoid extinction learning.

### Experiment 2: can bees solve a negative-patterning visual discrimination in a VR environment?

Bees (n = 20) were simultaneously trained to choose the single-colored gratings rewarded (A+ and B+) but not the compound grating (AB-) (Fig. 3). Conditioning consisted of three consecutive phases totalizing 32 trials (two phases of 11 trials and one of 10 trials in a random sequence; see Table S2) during which either A or B or AB was presented alone. At the beginning of each conditioning trial, the stimulus was displayed to the right or left (± 50° from the body axis) following a pseudo random sequence (10-trial phase: L, R, R, L, R, L, L, R, L, R; 11-trial phase: L, R, R, L, R, L, L, R, L, R, R). The single-colored gratings (CS+) were rewarded with sucrose solution when they were centered on the screen by the bee (alignment with the body axis), while the composite grating was not rewarded under the same conditions (CS-). Centered stimuli remained blocked in this position during 8 s, to allow sucrose delivery in the case of the CS+. In the case of the CS-, the stimulus also remained blocked during 8 s but without reward. Bees experienced 8 A+ trials, 8 B+ trials and 16 AB-trials, which allowed equating CS+ and CS-experiences. The same stimulus was never shown more than twice in a row (Table S2). Conditioning phases were separated by one hour during which bees rested on the miniature treadmill.

**Figure 3.**
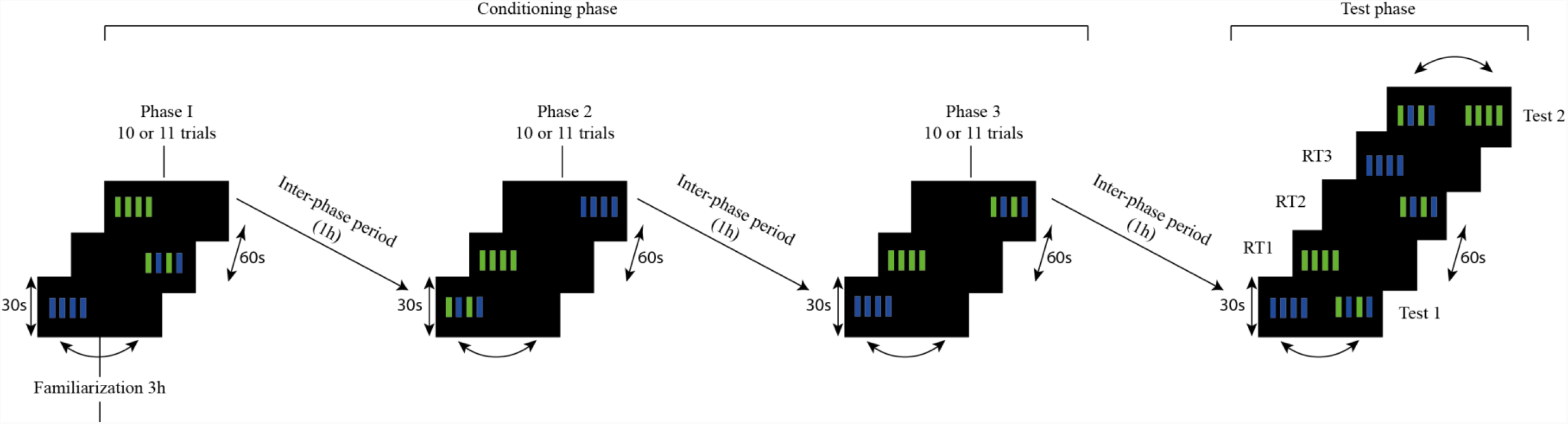
Experimental schedule of Experiment 2: negative patterning. Bees started with a familiarization phase with the setup in the absence of visual stimulation, which lasted 3 hours. The experiment consisted of 3 conditioning phases and a test phase. Each conditioning phase included either 10 or 11 trials during which either A, B or AB was presented alone on the right or on the left of the screen (± 50° from the body axis of the bee). A and B were rewarded with a sucrose solution (both CS+) when they were centered on the screen by the bee (0° from its body axis), while AB was not rewarded (CS-) under the same conditions. Trials lasted a maximum of 30 seconds. They were ended after reward delivery. The intertrial interval was 60 s. The presentation of A, B and AB was pseudo-randomized during the three conditioning phases (see Table S2). Each conditioning phase was separated by 1h. One hour after the last conditioning phase, bees were subjected to non-reinforced tests in which either CS+ was opposed to the CS-. Test sequence was randomized from bee to bee. Three refreshment trials were interspersed between tests to avoid extinction.

One hour after training, bees were subjected to two non-reinforced tests opposing the composite grating to each single-colored grating (i.e. AB vs A and AB vs B, Fig. 3, right). The relative position of the stimuli (right or left) was randomized from bee to bee. Centering the stimuli during these dual-choice situations did not result in blocking stimulus position in front of the bee. Thus, bees were able to switch from one stimulus to the other during the tests. Tests lasted 30 s and were separated by three refreshment trials (one per stimulus, A+, B+, AB−).

## Statistical analyses

For each bee and test situation, we recorded its first choice (first stimulation centered) and the time spent fixating each stimulus. In the case of the first variable, we calculated the proportion of bees first choosing the CS, the novel stimulus (NS) or not making any choice (NC) during the test. Data were bootstrapped to represent these proportions with their 95% confidence interval. To compare them, we used generalized linear mixed models (GLMMs) in a binomial family. For each model, the subjects were considered as a random factor to account for the repetitive measurement design. The times spent fixating each stimulus were compared using a Wilcoxon U rank test. All statistical analyses were done using the R 3.2.3 software (R Development Core team, 2018). Packages lme4 was used for GLMMs.

## Results

### Experiment 1: do bees treat a visual compound as the sum of its components?

#### Training with a single-colored grating

Bees trained with the green or the blue grating (A+, B+) did not differ in their test performances when these were quantified in terms of their first choice (GLMM; *A vs. B:* Group*Choice effect: χ^2^=2.37, df:2, *P*=0.30, NS; *A/B vs AB*: Group*Choice effect: χ^2^=4.40, df:2, *P*=0.11, NS) and stimulus-fixation time (Mann-Whitney; *A vs. B*: U=200, *P*=1, NS; *A/B vs AB:* U=144, *P*=0.13, NS). Performances were, therefore, pooled and expressed in terms of CS choice (Fig 4a) or fixation time (Fig. 5a), irrespective of the grating (blue or green) trained.

**Figure 4.**
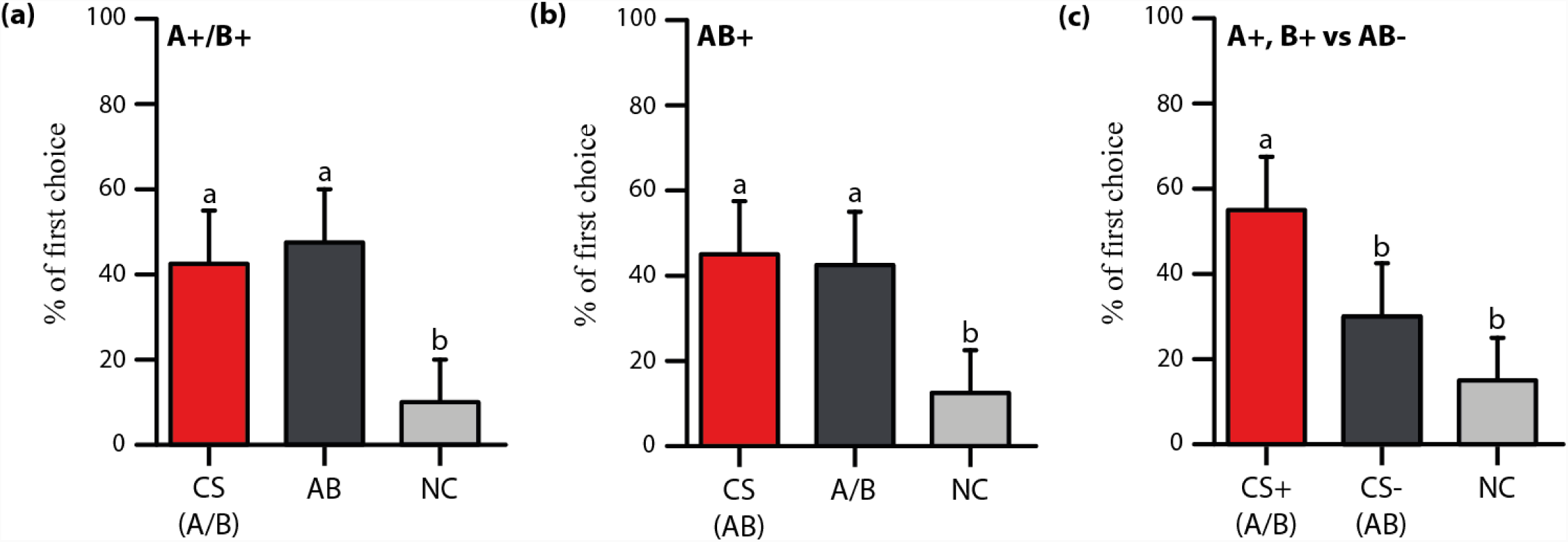
First choice of bees during the tests following absolute elemental conditioning (a,b) or negative-patterning conditioning (c). **(a)** Percentage of bees (n=40 bees, 20 for A+ training and 20 for B+ training) (± 95% confidence interval) first choosing the CS (A/B), the alternative stimulus AB or not making any choice (NC) during the test in which A or B were opposed to AB. The CS was either A or B. The CS bar represents the pooled performance corresponding to the presentation of A and B when these were trained. **(b)** Percentage of bees (n=20 bees) (± 95% confidence interval) choosing first the CS (AB), the alternative stimuli A/B or not making any choice (NC) during the test in which AB was opposed to A or B. **(c)** Percentage of bees (n=20 bees) (± 95% confidence interval) first choosing the CS+ (A/B), the CS-(AB) or not making any choice (NC) during the test in which A and B were opposed to AB. The CS+ bar represents the pooled performance corresponding to the presentation of A and B. Different lower-case letters indicate significant differences (*P<*0.05).

**Figure 5.**
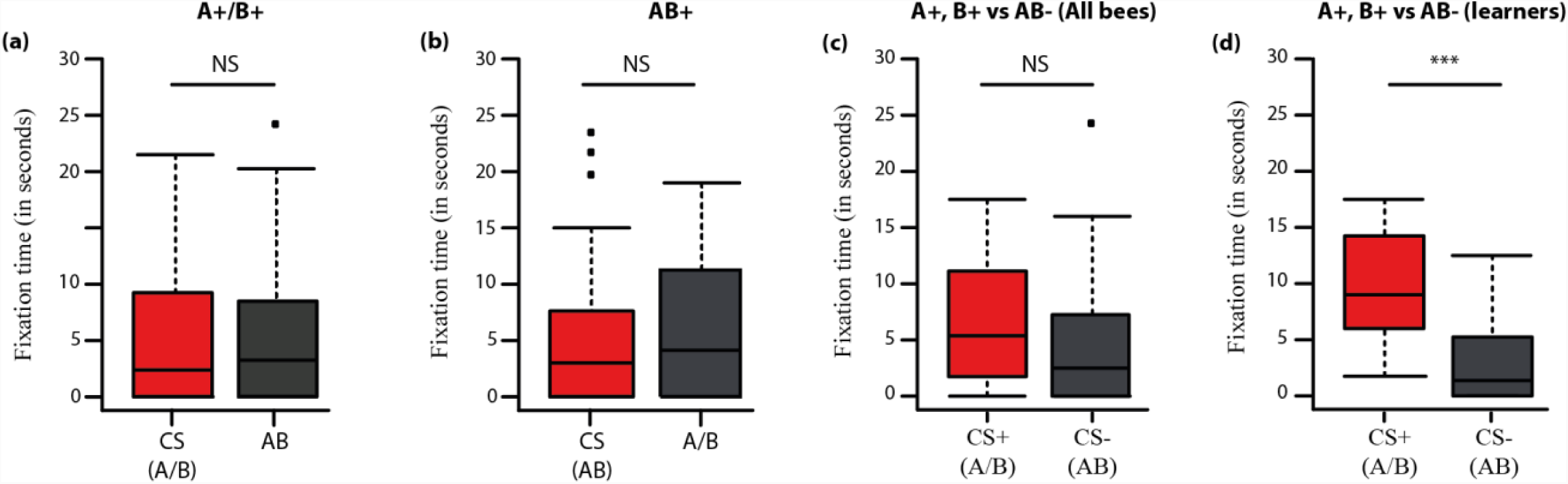
Fixation time of bees during the tests following absolute elemental conditioning (a,b) or negative-patterning conditioning (c,d). **(a)** Fixation time of the CS (A+/B+ pooled; red boxplot) and of the alternative stimulus AB (grey boxplot) during the test in which A and B were opposed to AB (n=40 bees, 20 for A+ training and 20 for B+ training). **(b)** Fixation time of the CS (AB+; red boxplot) and of A/B (times pooled; grey boxplot) during the test in which AB was opposed to A and B (n=20 bees). **(c)** Fixation time of the CS+ (A+/B+ pooled; red boxplot) and of the CS-(AB-; grey boxplot) during the test in which A and B were opposed to AB (n=20 bees). **(d)** Fixation time of the CS+ (A+/B+ pooled; red boxplot) and of the CS-(AB-; grey boxplot) during the test in which A and B were opposed to AB in the case of bees having chosen correctly (first choice for the CS+) during the test (n=17). In all cases the fixation time is expressed in s (median, quartiles and outliers). *: *P<*0.05, **: *P<*0.001, ***: *P<*0.0001, NS: non-significant.

Bees trained with a single-colored grating preferred their CS to the novel single-colored grating (test A vs B, Fig. S2a). The proportion of bees first choosing the CS was significantly higher than that of bees first choosing the novel single-grating (CS: 67.5 %; NS: 22.5 %) or not making any choice (NC: 10 %) (Fig. S2a, GLMM: CS vs. NS: z_357_=−3.68, *P*< 0.001; CS vs. NC: z_357_=−4.68, *P*<0.0001; NS vs. NC: z_357_=−1.48, *P*=0.14, NS). Thus, bees learned both trained single-colored gratings and discriminated them efficiently from each other. In consequence, they spent significantly more time fixating the former than the latter during the test (Fig S2b, Wilcoxon test: U=548, *P*<0.001).

When bees trained with the single-colored grating experienced this stimulus against the composite grating in a test (A or B vs. AB, Fig. 4a), they generalized their choice towards the compound grating and chose it at the same level as their CS: no significant preference for the CS was observed (CS: 42.5 %, NS: 47.5 %; z_357_=0.45, *P*=0.65, NS). The proportion of bees that did not make a choice remained significantly lower (NC: 10 %; CS vs NC: z_357_=−3.07, *P*<0.01; AB vs NC: z_357_=−3.41, *P*<0.001). Accordingly, bees did not spend significantly more time fixating the CS than the compound grating (Fig. 5a; U=285.5, *P*=0.46, NS).

Taken together, these results show that bees trained with a single-colored grating learned efficiently their CS in VR conditions and chose it preferentially against the alternative single-colored grating. They generalized their choice towards the composite grating, showing that they perceived the elemental gratings within it, consistently with a linear processing.

#### Training with a compound grating

Bees trained with the compound grating as CS performed similarly when this stimulus was confronted with either single colored grating (CS vs. A or CS vs. B; 1^st^ choice: Test*Choice: χ^2^=3.69, *P*=0.72, NS; fixation time: U=159, *P*=0.27, NS). The results were, therefore, pooled for analysis and graphical display. Bees trained with the compound grating chose equally the CS and the single-colored grating, be it A or B (Fig. 4b; CS: 45%; A/B: 42.5%; z_357_=−0.67, *P*=0.50, NS). The proportion of bees that did not make any choice (NC, 5%) remained significantly lower (CS vs NC: z_357_=−3.22, *P*<0.01; A/B vs. NC: z_357_=−2.67, *P*<0.01). Accordingly, bees spent the same time fixating the CS and the single-colored gratings (Fig. 5b; U=336, *P*=0.74, NS).

These results show that bees trained to a compound grating perceived the two grating components within the compound and generalized their choice towards the components, consistently with an elemental processing of the compound. This result confirms the findings obtained after conditioning with single-colored gratings and demonstrates that the processing inculcated by an elemental absolute conditioning is linear given that a compound is treated as the sum of its components. Thus, it is possible to ask if bees trained under a negative-patterning regime are able to inhibit the linear processing of a compound AB to solve this non-elemental discrimination.

### Experiment 2: can bees solve a negative-patterning visual discrimination in a VR environment?

There were no significant differences in performance regarding the first choice between both tests (Test*Choice: χ^2^=5.68, df=2, p=0.06), thus allowing the pooling of data in terms of CS+ vs. CS-responses. In these tests, the proportion of bees first choosing the CS+ (55%) was significantly higher than the proportions of bees choosing the CS-(30%) or not making any choice (15%) (Fig. 4c; CS+ vs. CS-: z_238_=−2.23, *P*<0.05; CS+ vs. NC: z_238_=3.55, *P*<0.001, CS-vs NC: z_238_=−1.58, *P*=0.11, NS). Thus, bees were able to solve the negative patterning discrimination as they suppressed linear responding to the compound.

The fixation time did not vary significantly between the CS+ and the CS-(Fig. 5c; U=44.5, *P*=0.13, NS), a result that may have been due to the relatively high proportion of non-learners in this experiment (30% CS-choosers and 15% non-choosers; see above). Restricting the analysis of the fixation time to the learners (i.e. bees that chose firstly the CS+) revealed a significant difference in the fixation time in favor of the CS+ (Fig. 5d; U=21, *P*<0.001). Yet, in this case, fixation time differed between single-colored gratings (W=108, *P*=0.01): bees spent significantly more time fixating the CS+ when it was the green grating (Fig. S3a; A+ vs AB-: U=75.5, *P*<0.001) than when it was the blue grating (Fig. S3b; B+ vs. AB-: U=175.5, *P*=0.51, NS).

From the 20 bees trained in the negative patterning task, five bees were successful as they first chose the CS+ and fixated it longer in both tests. Twelve bees were successful in only one of the tests. The remaining three bees were unsuccessful in both tests.

These results thus show that bees can solve a negative-patterning discrimination in the visual domain and in VR conditions. They responded more to the single-colored gratings and inhibited their otherwise lineal processing of the compound grating. In terms of fixation time, discriminating the green grating from the compound grating was easier than discriminating the blue grating from the compound grating.

## Discussion

Our results show for the first time that tethered honey bees walking stationary on a treadmill and trained with visual stimuli in a virtual environment learn a configural discrimination, the negative patterning. This task is considered a higher-order form of associative learning because elemental associative links between single stimuli and reinforcement (or absence of reinforcement) cannot account for solving this discrimination problem. The task is non-linear (the compound has to be treated as being different from the sum of its parts) and ambiguous (each single stimulus is as often reinforced as non-reinforced) ^3, 5^. The fact that bees learn the negative patterning under VR conditions shows that besides being able to solve patterning tasks in the olfactory domain^21-23^, they also master them in the visual domain. It also shows that the visual environment provided was sufficiently immersive and realistic despite the constraints imposed to the bees like the tethering and the absence of a perfect update of stimulus appearance relative to the bee’s movements (e.g. no stimulus looming).

Honey bees, like mammals and contrary to other insect species^37-38^, learn negative patterning discriminations using non-elemental strategies^24-25, 27-28^. This result raises the question of why bees succeed in this kind of learning while other insect species do not^37^. A possible answer may reside in the architecture of specific centers in the bee brain, which may be particularly adapted to mediate configural learning in the bee. In particular, the mushroom bodies (MBs) of the honey bee may be fundamental to this end. MBs are higher-order associative brain structures, which in the bee are multimodal and allow the combination of information pertaining to different sensory modalities (e.g. olfactory, visual, mechanosensory, gustatory)^39-40^. This is different from other insect species where MBs are mostly unimodal or dominated by a single sensory modality (e.g. in the the fruit fly *Drosophila melanogaster*). This multimodality may explain why bees can solve a negative patterning both in the olfactory and in the visual domain. Yet, a fundamental architecture principle for solving this task may be the existence of inhibitory feedback neurons at the level of the MBs.

GABA immunoreactive feedback neurons (also termed protocerebral-calycal neurons or PCT neurons) are a major component of the honeybee mushroom body^41^. They exhibit learning-related plasticity^42^ and provide putative inhibitory input to Kenyon cells and the pedunculus extrinsic neuron, PE1. PCT neuron activity may account for suppression of compound responses in configural tasks such as negative patterning. Sparse coding by Kenyon cells at the level of the MBs would yield a reduced activation of PCT neurons upon single stimulus presentation (A+, B+) but a supra-threshold activation during compound presentation (AB−), thus enabling GABAergic inhibition to suppress linear processing based on summation of neural responses. Pharmacological blockade of PCT neurons upon olfactory patterning tasks yielded results consistent with this hypothesis: in this case, bees loss the capacity to solve patterning tasks but maintained the capacity to solve elemental discriminations. Similar results are expected for visual patterning tasks; as PCT neurons provide inhibitory feedback not only to the basal ring, the olfactory input region of the MBs, but also to the collar, the visual input region of the MBs, blockade of their activity should also impair negative-patterning solving in the visual domain. It can be therefore proposed that configural learning in multiple domains requires multimodal MBs and retrograde inhibition onto MB input signals disrupting linear summation.

The success of bees in our VR setup shows that this environment is suitable for studying elemental^33-35^ and non-elemental learning under controlled laboratory conditions. It has the advantage of reproducing these learning forms, which are typically exhibited by free-flying bees, in tethered bees walking stationary on a fixed point of space. This facilitates the coupling of behavioral experiments with invasive methods aiming at dissecting the neural bases of different learning forms in the bee. The preparation is perfectible as stimulus looming or receding upon forward or backward movements were not available in our display. Yet they can be easily incorporated and the prediction is that this will improve considerably learning success, even if it is well established that learning success in negative patterning is always lower than in elemental learning due to the difficulty of the configural task^25^. In this way, the study of honey bee visual learning, which has historically suffered from the drawback of not enabling a mechanistic analysis due to the use of free-flying honeybees, will make relevant progresses in an immediate future.

## Acknowledgements

Our work was supported by the Human Frontier Science Program (grant RGP0022/2014), the French National Research Agency (grant ANR-13-BSV4-0004-0), the French National Research Center (CNRS) and the University Paul Sabatier of Toulouse. Lucie Hotier provided help in terms of beekeeping activities. Special thanks are due to Sébastian Weber for his technical assistance in designing and implementing the software for visual stimulus control and for his regular support throughout this project.

## Author contributions

A.B., A.A.-W. and M.G. designed the experiments. The experiments were conducted by A.B. and L.L. A.B. and L.L. analyzed the results and prepared figures and tables. A.B., A.A.-W., and M.G. wrote the manuscript. All authors reviewed and approved the final manuscript.

